# TIGIT drives the immunosuppressive environment by downregulation of metalloproteinases MMP2 and MMP14 in perihilar cholangiocarcinoma

**DOI:** 10.1101/2023.08.14.553195

**Authors:** Lara Heij, Konrad Reichel, Willem de Koning, Jan Bednarsch, Xiuxiang Tan, Julia Campello Deierl, Marian Clahsen-van Groningen, Tarick Al-Masri, Ronald van Dam, Juan Garcia Vallejo, Florian Ulmer, Sven Lang, Tom Luedde, Flavio G. Rocha, Edgar Dahl, Danny Jonigk, Mark Kuehne, Shivan Sivakumar, Ulf Neumann

## Abstract

**Background:** Checkpoint blockade in cholangiocarcinoma (CCA) is promising; however, little is known about the response to treatment in CCA subtypes. In this study, we investigated the spatial immune environment in combination with checkpoint expression in perihilar CCA (pCCA).

**Materials & Methods:** The levels of checkpoint molecules (PD-1, PD-L1, PD-L2, LAG-3, ICOS, TIGIT, TIM-3, and CTLA-4), macrophages (CD68), and T cells (CD4 and CD8) were assessed by multiplex immunofluorescence (mIF) in 50 patients. We investigated the transcriptomic profile using the NanoString Cancer Progression Panel, and validation was performed by mIF on tissue sections from 24 patients.

**Results:** The expression of checkpoint molecules TIGIT, CTLA-4, and LAG-3 alone and in combination with other checkpoint molecules was more abundant in the Central Tumor (CT) and Invasive Margin (IM) than in peritumoral tissue (PT) (CD4 and CD8 TIGIT p<0.0001 for both CD4 and CD8 CTLA-4, p<0.0001 and p < 0.001, respectively, and CD8 LAG-3 p < 0.05). MMP2 and MMP14 were differentially expressed in patients with high TIGIT expression.

**Conclusion:** The immune environment in pCCA is characterized by the expression of multiple checkpoints, demonstrating the complexity of ICI treatment. High TIGIT expression drives an immunosuppressive environment by modulating the extracellular matrix. Future clinical trials in pCCA could consider TIGIT as a therapeutically relevant target for (combination) treatment.

## Introduction

Cholangiocarcinoma (CCA) is a liver cancer that originates from the bile ducts and is associated with high mortality rates(1). Depending on the location of the cancer; it is classified as an intrahepatic cholangiocarcinoma (iCCA), perihilar cholangiocarcinoma (pCCA) or a distal cholangiocarcinoma (dCCA). Response to treatment is often limited, and surgery offers the best chance of cure(2, 3, 4). To date, the three different subtypes of CCA have often been pooled in the same clinical studies, despite their different clinical courses.

pCCA tumors are surrounded by nerves and large blood vessels, which can infiltrate the tumor and therefore influence survival. Additionally, the presence of small nerves in the tumor microenvironment (TME) can predict better survival(5, 6, 7). It is likely that the composition of the TME directly affects immune cell spatiality and co-expression of checkpoint molecules(8).

Immune checkpoint inhibitors (ICI) are effective against several cancer types. Not all patients respond equally, and reliable biomarkers to predict the response to (immuno)therapy are lacking(9, 10, 11).

The TME hosts many different cell types, and crosstalk between cell clusters exist. Immune cell composition and distribution can either facilitate or limit tumor growth(12, 13, 14). Ligand expression blocks immune cells by expressing co-inhibitory receptors. Co-stimulatory and co-inhibitory checkpoints exist and play a role in antitumor responses (15). The co-expression of checkpoint molecules in immune cells correlates with survival in several cancer entities(16, 17). Combination treatment with immunotherapy and chemotherapy results in better overall survival in patients with biliary tract cancer(18).

In CCA, much is unknown about the TME, and this is an ongoing field of investigation. A key aspect of the TME is remodeling of the extracellular matrix (ECM) by cancer cells and fibroblasts, which can have various immunomodulatory functions(19). CD4 T cells, including Tregs, cytotoxic CD8 T cells, and macrophages, are major players within the immune landscape (8). However, little is known about its presence and checkpoint expression in pCCA. Existing cancer immunotherapeutics mostly consist of anti-PD-1, anti-PD-L1, and anti-CTLA-4 targeting antibodies; however, many more checkpoint antibodies are in development. In this study, we integrated the co-inhibitory checkpoints TIM-3, LAG-3, CTLA-4, TIGIT, PD-1, PD-L1, PD-L2, and the co-stimulatory checkpoint ICOS, all in various stages of clinical development. Additionally, we analyzed the presence of lineage markers CD4, CD8, and CD68 in samples of primary resected patients with pCCA using multiplexed imaging. Our aim was to characterize the spatial immune environment and expression of immune checkpoints (see Figure 1 for the study workflow).

**Figure 1.**
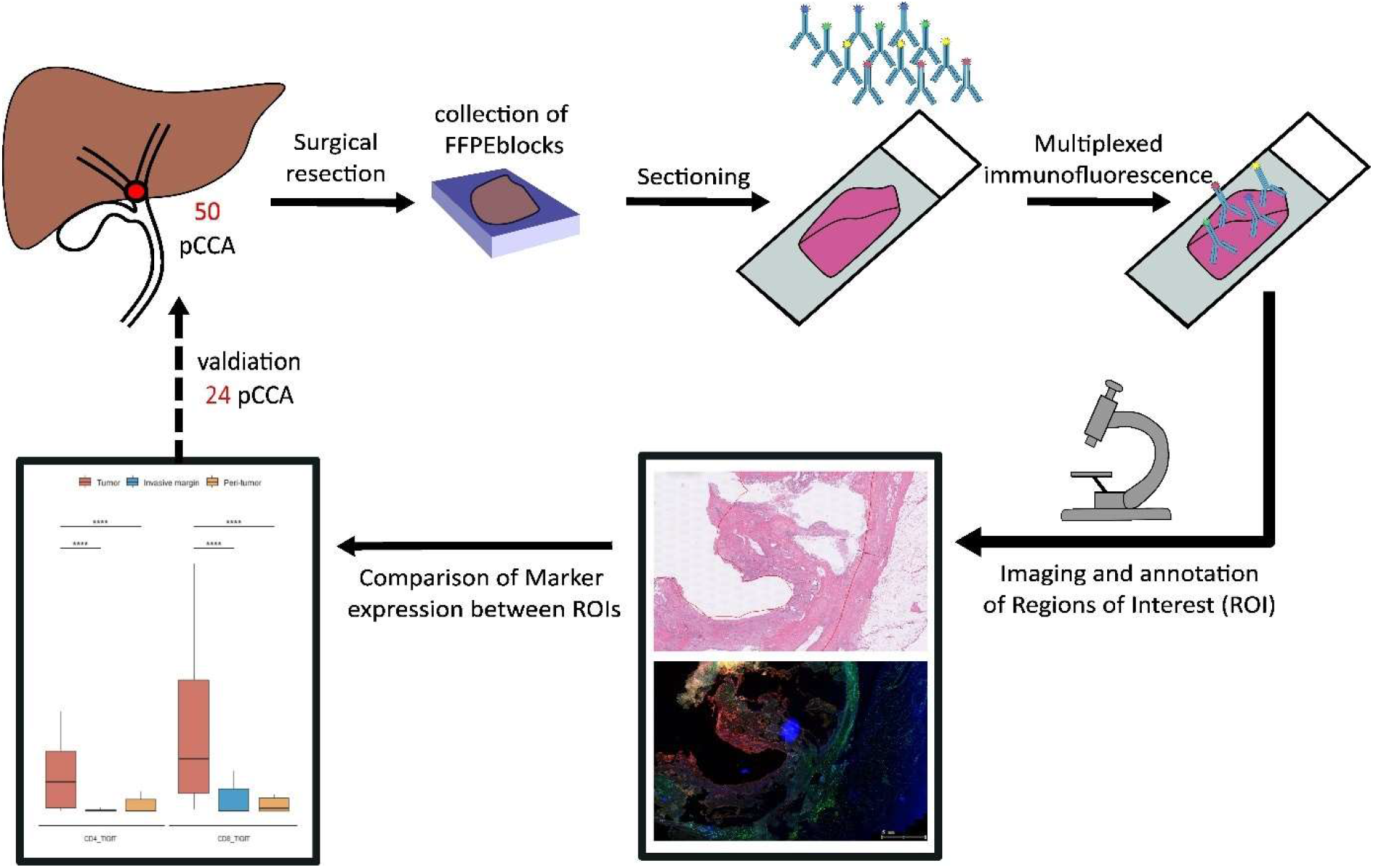
Overview of the study workflow. The study included 50 patients with pCCA who underwent surgical resection. A representative FFPE block was obtained from the pathology archive. The slides were cut and prepared for a multiplexed imaging workflow. First, DAPI staining was performed and then using the antibodies from all panels. The tissue slides were scanned and analyzed using Tissuegnostics Software. After quantification of the cells and measurement of co-expression, statistical analysis was performed. We validated our findings in an additional cohort of 24 patients using an adapted panel with a subset of the original markers. *pCCA: perihilar cholangiocarcinoma, FFPE: Formalin-fixed paraffin-embedded, DAPI: 4′,6-diamidino-2-fenylindool nuclear staining*.

## Materials/methods

### Patient Recruitment

FFPE blocks were collected from the pathology archive of 50 patients diagnosed with pCCA between 2010 and 2019 who were fit for surgery with curative intent; for specific patient characteristics, see Supplementary Table 1. The study was conducted in accordance with the requirements of the Institutional Review Board of RWTH-Aachen University (EK 106/18 and EK 360/19), Declaration of Helsinki, and Good Clinical Practice Guidelines (ICH-GCP).

### Sample Collection

Hematoxylin and Eosin (HE) slides were collected, and the slide with the vital tumor and presence of interface area was selected by a pathologist (LH). The selected block was used for further processing of the multiplexed imaging workflow using the Tissuefaxs method (TissueGnostics, Austria). For tumor staging and grading, TNM classification was performed according to the AJCC/UICC 8th edition.

### Whole slide multiplexed immunofluorescence (mIF)

All FFPE samples were subjected to multiplexed immunofluorescence (mIF) in serial 5 μm histological tumor sections obtained from representative FFPE tumor blocks. FFPE blocks were carefully selected in the presence of the tumor region and, if available, invasive margin and peritumoral tissue. The sections were labeled using the Opal 7-Color fIHC Kit (PerkinElmer, Waltham, MA, USA). Antibody fluorophores were grouped into a panel of five antibodies. The order of antibody staining was kept constant on all sections, and sections were counterstained with DAPI (Vector Laboratories). The multiplexed immunofluorescence panel consisted of CD4, CD8, CD68, PD-1, PD-L1, PD-L2, ICOS, TIGIT, TIM-3, CTLA-4, and LAG-3 (Supplementary Table 2). All antibodies were diluted with an Antibody Diluent (Background Reducing Components, Dako, Germany). Secondary antibodies were applied using the ImmPRESS™ HRP (Peroxidase) Polymer Detection Kit (Vector Laboratories, US). TSA reagents were diluted with 1× Plus Amplification Diluent (PerkinElmer/Akoya Biosciences, US).

The manual for mIF was described by Edwin R. Parra’s protocol(20), in which the first marker was incubated after FFPE sections were deparaffinized in xylene and rehydrated in graded alcohols. The second marker was applied the following day. The third marker was applied on the third day, which was continued for all markers. After the five sequential reactions, the sections were cover-slipped with VECTRASHIELD® HardSet™ Antifade Mounting Medium.

The slides were digitally scanned using the TissueFAXS PLUS system (TissueGnostics, Austria). Image analysis was performed in different regions of interest (ROI) in each image (only if present on the slide): central tumor (CT), invasive margin (IM), and peritumoral areas (PT). The size of the ROI varied for each slide.

Strataquest software was used to analyze antibody staining, and cell counts. Library information was used to associate each fluorochrome component with an mIF marker. All immune cell populations were quantified as positive cells per mm^2^ using cell segmentation, and thresholds were set manually under the supervision of a pathologist (LH). Positive cell counts were categorized based on thresholds, and values above the threshold were considered positive. Immune cell expression was calculated as a percentage throughout the project. The checks were performed by a pathologist (LH).

### Statistical analysis

The primary endpoint of this study was to compare the immune cell composition, distribution, and co-expression of checkpoints in patients with pCCA. Group comparisons between ROIs were performed using CT, IM, and PT. The different ROIs were compared within the entity itself with the aim of visualizing spatial differences within the same slide. Associations between immune cell subsets and clinical variables were investigated using binary logistic regression. Therefore, immune cell expression data were converted into a dummy variable (high expression vs. low expression) using median expression as a cut-off for grouping between high and low expression.

Group comparisons were conducted using the Mann-Whitney U test or t-test for continuous variables, while the χ2 test and Fisher’s exact test in accordance with scale and number count were used for categorical variables. The Wilcoxon-matched-pairs-test or the paired t-test was applied to determine statistically significant differences between the values of immune cells within the ROIs. The level of significance was set at α = 0.05, and p-values were calculated 2-sided testing. All statistical calculations were performed using Python (v25, IBM, Armonk, NY, USA).

### Nanostring Pan Cancer Progression Panel

RNA was extracted from the IM and PT subregions, and three consecutive 20-µm sections were obtained from each FFPE block. The sections were then immediately transferred to sterile microcentrifuge tubes and stored at room temperature. Microtome blades were replaced, and between blocks, the equipment was sterilized with RNase AWAY (Life Technologies, Carlsbad, CA, USA). Xylene deparaffinization and RNA isolation were performed according to the manufacturer’s instructions using the RNeasy FFPE Kit. RNA concentration and subsequently, both quality and quantity were measured using a NanoDrop 2000 spectrophotometer (Thermo Fisher Scientific, Waltham, MA) and Bioanalyzer (Agilent, Santa Clara, CA).

The NanoString nCounter Reporter CodeSet (NanoString Technologies, USA) was used in this study. This panel included 770 genes covering a range of cancer-related pathways, including angiogenesis, extracellular matrix (ECM) remodeling, epithelial-to-mesenchymal transition, and metastasis (Supplementary Table 3).

Statistical testing and data visualization were performed using R Statistical Software (v.4.1.2) (R Core Team, 2022). We used the R packages ggplot2 (Wickham, 2016) and EnhancedVolcano (Blighe et al., 2021) for data visualization. Heatmaps were generated using the Log2-normalized count data of significantly differentially expressed genes (|Log2 Fold change| > 0.5, p-value < 0.05). Genes determined to be outliers were removed using Tukey’s rule were removed (Tukey, 1977). The heatmap was visualized using the web-based tool Morpheus by the Broad Institute (RRID: SCR_017386).

Pathway enrichment analyses were performed with differentially expressed genes (|Log2 fold change| > 0.5, p-value < 0.05) using Metascape (Y. Zhou et al., 2019) and ClueGo (Bindea et al., 2009).

### Validation cohort

We then selected FFPE blocks from 24 patients diagnosed with pCCA between 2020 and 2022, who were fit for surgery with curative intent. The validation panel included CD4, CD8, TIGIT, CTLA-4, and LAG-3. The slides were digitized, and outlining was performed (Supplementary Table 4 for an overview of patient characteristics).

## Results

For this study, ROIs were analyzed from 50 patient samples, the outlined ROIs demonstrated the CT, IM, and PT areas, but not all areas were available for every selected slide. The outlining was performed by a pathologist (LH). Positive cell counting was performed and measured as percentage per mm^2^.

### CTLA-4, TIGIT, ICOS and LAG-3 are abundant in the immunosuppressive environment

In this study, we examined checkpoint expression in CD4+ and CD8+ T cells and CD68+ macrophages. There were clear differences in the spatial distribution of checkpoint marker expression between the ROIs (Figure 2A-D). In addition to the difference in the levels of checkpoint-expressing immune cells, the levels of total CD8+ T cells and CD68+ macrophages were also elevated in CT compared to PT (p=0.05 and p <0.001, respectively). This was not the case for CD4+ cells, which were present in similar numbers across ROIs (Figure 2 E).

**Figure 2.**
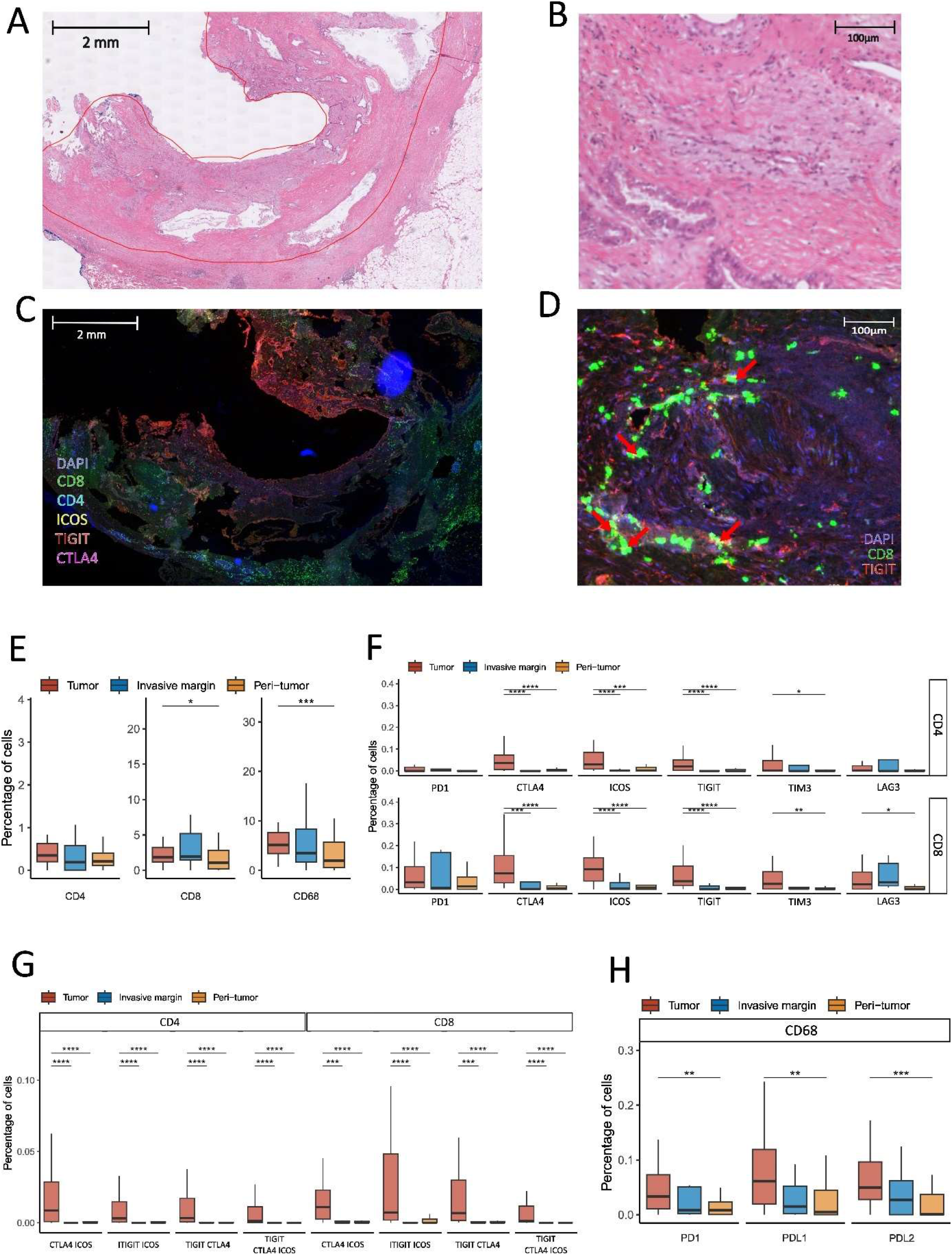
Checkpoint marker expression between and within the different ROIs in pCCA. Fifty tissue slides of pCCA were analyzed using multiplexed immunofluorescence. Markers focused on immune checkpoints included lineage markers for CD4+ and CD8+ T cells and CD68+ macrophages. Annotation of the region of interest (ROI) into the central tumor, invasive margin, and peri-tumor was performed by a pathologist (LH), and the percentage of positive cells for the given markers was compared between ROIs. **A:** Representative image of hematoxylin and eosin (HE) staining is shown, where the tumor ROI is outlined in red. **B:** Close-up view of the area within the tumor region. **C:** Multiplexed immunofluorescence image using one of the panels of the same region, as shown in A. **D:** TIGIT expression in CD8 cells is shown for the same representative field of view as in B. Red arrows indicate examples of CD8 and TIGIT co-expression. **E:** Differences in the abundance of lineage marker-positive cells between the regions of interest. **F:** Expression of checkpoint markers in CD4 and CD8 positive cells is clearly elevated in the tumor. **F:** Co-expression of checkpoints ICOS, CTLA4, and TIGIT in CD4 and CD8 positive cells. **H:** Checkpoint expression in CD68 positive cells **E-H**: P values were generated using the Mann-Whitney U test. * p < 0.05, **p < 0.01, ***: p < 0.001, **** p < 0.0001. *pCCA: perihilar Cholangiocarcinoma, ROI: Region of Interest, HE: Hematoxylin and Eosin, ICOS: Inducible T-cell costimulatory, CTLA4: Cytotoxic T-lymphocyte-associated protein 4, TIGIT: T cell immunoreceptor with Ig and ITIM domains*.

Looking at the expression of checkpoints in the T cell populations, we observed an increase in most of the assessed markers on CT. In the CD4+ and CD8+ compartments, TIGIT, CTLA-4, and ICOS expression was clearly increased compared to both the IM and PT (for CD4 and CD8 TIGIT p=<0.001 for both ROIs, for CD4 CTLA-4, p=<0.001 for both ROIs and CD8 CTLA-4 CT-IM p=<0.001 and CD8 CTLA-4 CT-PT p=<0.0001 and CD4 ICOS CT-IM p=<0.0001 and CT-PT p=<0.001 and for CD8 ICOS p=<0.0001 for both ROIs). TIM-3 was also elevated in the CT for both cell populations, but only when compared to the PT (CD4 TIM-3, p<0.05; CD8 TIM-3, p<0.01). There was an increase in LAG-3 positive cells for CD8 and CD4 in CT compared to PT, but this only reached significance for CD8 cells (p<0.05). Surprisingly, we did not identify any statistical differences in PD-1 expression between the ROIs (Figure 2F). Furthermore, we were able to show increased co-expression of different combinations of TIGIT, CTLA-4, and ICOS in the tumor (Figure 2G). Although TIGIT, CTLA-4, and ICOS were more clearly elevated than other checkpoints, PD-1, TIM-3, and LAG-3 also contributed to the co-expression of checkpoints (Supplementary Table 5).

Within the population of CD68+ macrophages, we observed an increase in PD-1, PD-L1, and PD-L2 within the CT compared to the PT (Figure 2H) (p=0.01, p <0.01, p<0.001, respectively) but did not observe any significant increase in checkpoint co-expression between the ROIs.

### Validation of the checkpoint distribution

We then validated the findings in a second cohort; here, we included 24 patients between 2020 and 2022 (see Supplementary Table 4 for patient characteristics). For validation, we included significant markers in relation to TIGIT. CD4, CD8, TIGIT, CTLA-4, and LAG-3 were included using the same methods as previously described. PT, CT, and IM were outlined by a pathologist (LH) (see Supplementary Table 6) for an overview of the ROIs.

In support of the first cohort, we uncovered high levels of TIGIT expression within the CT (for CD4 TIGIT, p<0.01; CD8 TIGIT, p<0.001). The number of CD4 and CD8 cells expressing TIGIT and CTLA-4 was higher in the tumor than in the PT. Despite this, the increase in CTLA-4 did not reach significance in CD4 cells, and CD8 CTLA-4 was increased in CT (p<0.05). For LAG-3, we observed a significant increase in positive cells in the IM compared to the other two ROIs (CT-IM, p<0.001; IM-PT, p<0.05). In addition, we observed LAG-3 co-expression with CTLA-4 in CD8 cell tumors (p<0.05) and CD4 TIGIT combined with LAG-3 (p<0.01). There was also a small percentage of CD4 and CD8 cells co-expressing CTLA-4, LAG-3, and TIGIT in the CT group (for CD4 p<0.01, and for CD8 p<0.001) (Figure 3A).

**Figure 3.**
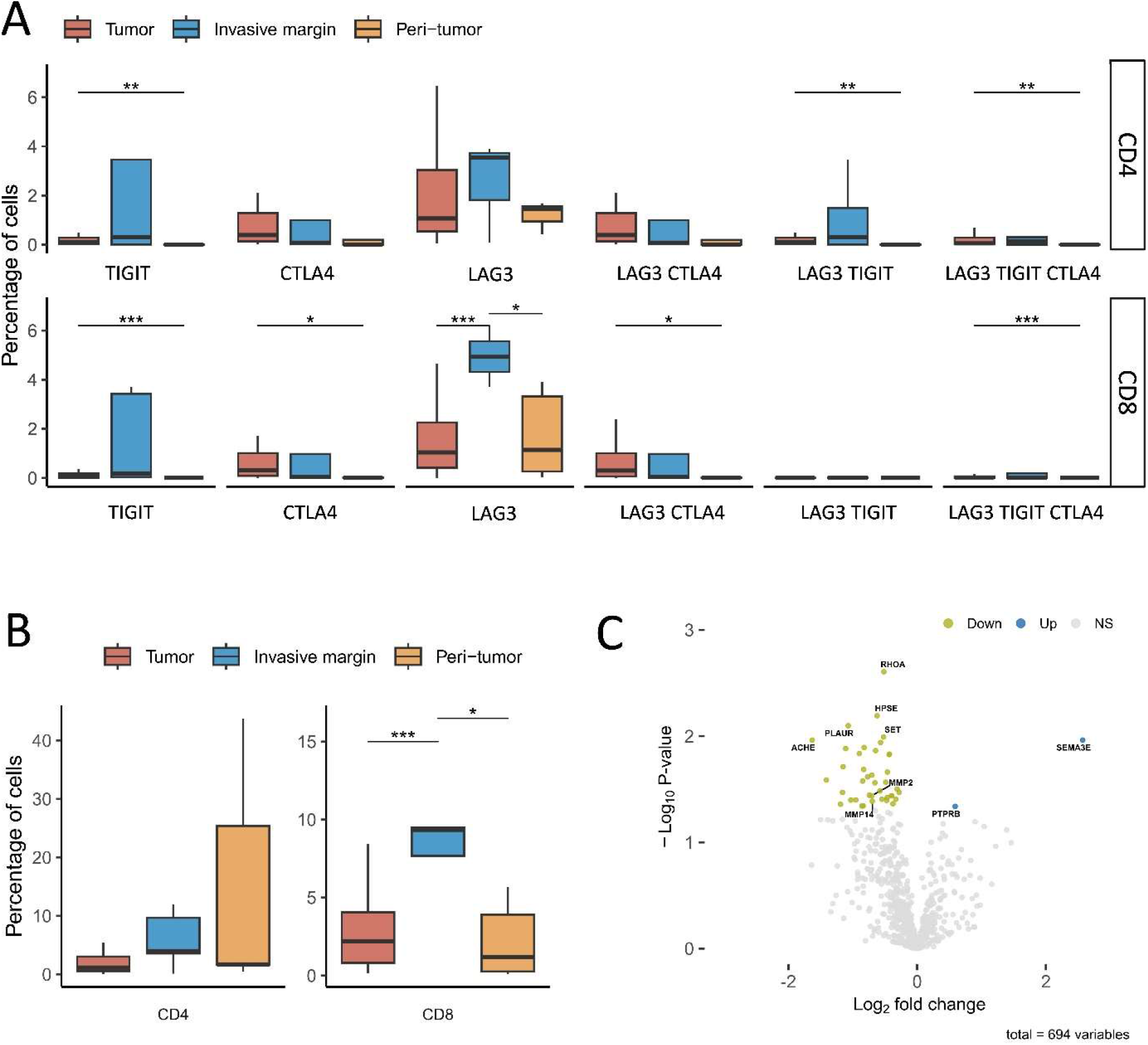
Validation of checkpoint co-expression and transcriptional analysis of TIGIT high tumors. For validation 24 tissue slides of pCCA were analyzed using multiplexed immunofluorescence. The panel was adapted to focus on key checkpoints in CD8+ and CD4+ T cells. Annotation of region of interests (ROI) into T, IM and PT was done by a pathologist as done for the first cohort percentage of positive cells for given markers was compared between ROIs. **A:** Differences in co-expression of checkpoint markers in CD4 and CD8 positive cells. **B:** CD4 and CD8 maker expression across the ROIs. **A-B:** p-values were generated using Mann-Whitney U test. * p < 0.05, **p < 0.01, ***: p < 0.001, **** p < 0.0001. **C:** Based on the results from the multiplexed images we chose 5 patients with low TIGIT levels and 6 patients with high TIGIT levels and extracted RNA from the Tumor and IM subregions for targeted transcriptomic analysis using the NanoString PanCancer Progression Panel. Differential gene analysis between the two groups was performed. Significant down and upregulated genes are shown in green and blue respectively. Selection of significant genes was based on a Log2 Fold change cutoff of > 0.5 and p-value cutoff of < 0.05. *ROI: Region of interest, CT: Central tumor, IM: invasive margin, PT: peri-tumor, TIGIT: T cell immunoreceptor with Ig and ITIM domains, RNA: Ribonucleic acid*.

Comparable to earlier results, CD4 cell levels were not significantly enriched in any of the ROIs, but CD8 levels were elevated in IM compared to both the PT and the tumor region (CD8 CT-IM p<0.001 and CD8 IM-PT p<0.05) (Figure 3B).

### Nanostring PanCancer Transcriptome analysis reveals downregulation of Metalloproteinases

Based on the previous analysis, we selected six patients with high TIGIT levels and five patients with low TIGIT levels for transcriptome analysis using the NanoString PanCancer Progression Panel. A total of 28 genes were significantly downregulated and 2 significantly upregulated in patients with high levels of TIGIT (Figure 3C and supplementary table 7). Among the downregulated genes were MMP2 and MMP14. Functional enrichment of downregulated genes revealed the enrichment of genes associated with collagen degradation in downregulated genes.

## Discussion

The diversity between cholangiocarcinoma subtypes is high, and therapeutic targets are evolving. However, not all patients respond equally to these regimens. In iCCA, genetic mutations are known to play a role in tumor heterogeneity, and molecular testing is useful in the case of cancer progression (11). Different spatial immunophenotypes have been described in iCCA, and the inflamed phenotype is correlated with better overall survival(21). This phenotype, with many T cells in the cancer environment, has been demonstrated in 11% of iCCA cases(8). CD4 T regulatory cells are present in an immunosuppressive environment and are indicative of poor outcome(9, 22). CD8 cytotoxic cells facilitate an antitumor response and are more abundant in patients with better outcomes (23, 24). In addition to the presence of specific subtypes of immune cells, their activation status(25, 26) and expression of checkpoint molecules are known to play an important role.

In this study, we investigated the expression of checkpoint molecules in patients with pCCA and provided an overview of their spatial distribution. Our results have yielded several findings. The expression of TIGIT alone and in combination with other checkpoint molecules was more abundant in CT and IM than in PT. TIGIT interacts with CD155 expressed on cancer cells or antigen-presenting cells and exerts an immunosuppressive effect(27). This finding could indicate a possible effect of TIGIT antibody therapy in combination with other therapeutic agents. In addition, LAG-3 and CTLA-4 were more abundant in patients with pCCA. LAG-3 is a co-inhibitory molecule that is believed to have a suppressive effect on CD8+ T cells. LAG-3 and CTLA-4 are therapeutic targets in advanced clinical trials(28, 29, 30)

Blocking TIGIT with monoclonal antibodies and a combination of TIGIT and PD-1 blockade in particular prevented tumor progression, distant metastasis, and tumor recurrence in in vivo models(28). In Hepatocellular carcinoma (HCC), the expression of TIGIT is indicative of an immune-exhausted phenotype, suggesting that patients with TIGIT expression might not be as sensitive to immunotherapy. Our results support this finding, since high levels of ICOS and the expression of multiple checkpoint molecules on T cells are factors that negatively influence the host immune response. This is in line with the results of the TOPAZ trial, where subgroup analysis demonstrated that PD-L1 blockade therapy was more effective in iCCA.

In addition, LAG-3 is a therapeutic target, and results in melanoma have been promising. Previous studies have demonstrated that LAG-3 is upregulated in various types of tumors and suppresses the proliferation, activation, and effector functions of T-cells.

CTLA-4 primarily works by outcompeting the co-stimulatory receptor CD28 for CD80/CD86 binding, and therefore removes vital co-stimulatory signals required for an efficient T cell response. A combination of PD1 and CTLA4 checkpoints is commonly used in other cancer types and is thought to synergize well(31). Supporting its immunosuppressive role in the TME, iCCA patients with high levels of CTLA-4 have been shown to have a worse prognosis (32), and a study suggests a beneficial response in a subset of iCCA patients(33). Here, we showed high levels of CTLA in pCCA tumors, implying that pCCA could benefit from a similar treatment regimen.

IM is where the cancer is the most aggressive and invades the surrounding tissue. The presence of effective immune cells in the surrounding regions of the tumor is an important finding since immune cells can migrate to the IM. Further research is needed to investigate whether immune cells can be recruited to the IM when they are released by ICI therapy. The zonation in the distribution of checkpoint molecules could be of importance when expression of the checkpoints is determined on a biopsy and used as a biomarker to enrol patients for therapy.

Furthermore, we identified downregulation of matrix metalloproteinases MMP2 and MMP14 in patients with high TIGIT levels. MMP2 and MMP14 can break down the extracellular matrix (ECM) by cleaving collagen. Collagen production is commonly induced by cancer cells, is associated with an unfavorable patient outcome, and is known to promote a TME characterized by the exclusion of effector T cells (34). In addition to restricted infiltration, high collagen density was also shown to modulate T cell activity from a cytotoxic to a more regulatory state(35). CD8 cells can directly recognize collagen via the LAIR1 receptor, leading to T cell exhaustion, which has been suggested as a mechanism for immunotherapy resistance in human lung tumors(36). Together, this suggests that remodulation of the ECM by decreasing collagen degradation might be connected to the observed upregulation of immunosuppressive molecules. On the other hand, MMPs, such as MMP2 and MMP14, are commonly upregulated by cancer cells and are also involved in other aspects of the TME, such as cancer cell migration and angiogenesis, making their role more complex (37, 38).

The limitations of the present study are that we only included treatment-naïve patients; we do not know anything about the potential change in the immune environment after therapy. The NanoString Cancer Progression panel includes a limited number of genes, and potentially a phenotype driven by genes not included in the NanoString panel would not be detected by this assay.

The expression of multiple checkpoints potentially indicates that pCCA will be PD-1 blockade refractory when treated with PD-1 blockade therapy alone. Targeting pCCA could indicate that a combination of different immune checkpoint agents is a better approach than PD-1 blockade monotherapy. It is important that these new therapeutic strategies are tested, potentially in combination immunotherapy within a chemotherapy setting.

## Abbreviations

CCA: Cholangiocarcinoma
CTLA-4: Cytotoxic T-lymphocyte-associated protein 4 D
API: 4′,6-diamidino-2-fenylindool
DCCA: Distal cholangiocarcinoma
ECM: Extracellular matrix
FFPE: Formalin-fixed paraffin-embedded
HCC: Hepatocellular carcinoma
HE: Hematoxylin and Eosin
ICCA: Intrahepatic cholangiocarcinoma
ICI: Immune checkpoint inhibitor
ICOS: Inducible T-cell costimulatory
IM: Invasive margin
LAG-3: Lymphocyte-activation gene 3
mIF: Multiplexed immunofluorescence
MMP2: Metalloproteinase-2
MMP14: Metalloproteinase-14
PCCA: perihilar cholangiocarcinoma
PD-1: Programmed death 1
PD-L1: Programmed death-ligand 1
PD-L2: Programmed death 1 ligand 2
PT: Peritumoral area
RNA: Ribonucleic acid
ROI: Region of interest
RWTH: Rheinisch-Westfälische Technische Hochschule
TIGIT: T cell immunoreceptor with Ig and ITIM domains
TIM-3: T cell immunoglobulin and mucin-domain containing 3
TME: Tumor microenvironment
Treg: T regulatory cell
CT: Central tumor

