## Supplementary files for "TIGIT drives the immunosuppressive environment by downregulation of metalloproteinases MMP2 and MMP14 in perihilar cholangiocarcinoma"

Supplementary table 1. Patient characteristics

| | Total | Percentage |
|---|---|---|
| **Included pCCA patients** | **50** | |
| **Mean age (years)** | 64.32 | |
| **Mean overall survival (months)** | 47.96 | |
| **Gender** | | |
| %Male | 30 | 60% |
| %Female | 20 | 40% |
| **Tumor stadium (T)** | | |
| %UICC T1 | 0 | 0% |
| %UICC T2 | 31 | 62% |
| %UICC T3 | 13 | 26% |
| %UICC T4 | 6 | 12% |
| **Nodal status (N)** | | |
| %N0 | 22 | 44% |
| %N1+N2 | **28** | 56% |
| **Tumor grading (G)** | | |
| %G1 | 0 | 0% |
| %G2 | 37 | 74% |
| %G3 | 9 | 18% |
| %G4 | 1 | 2% |
| %not known | 3 | 6% |
| **Perineural Invasion (Pn)** | | |
| %Pn0 | 5 | 10% |
| %Pn1 | 36 | 72% |
| %not known | 9 | 18% |
| **Residual Tumor (R)** | | |
| %R0 | 8 | 16% |
| %R1 | 42 | 84% |
| %not known | 0 | 0% |
| **Lymphovascular invasion (L)** | | |
| %L0 | 34 | 68% |
| %L1 | 13 | 28% |
| %not known | 3 | 6% |

, ,

Supplementary Table 2. Antibody panels

| Antibody | Marker | Dilution | Incubation time | Manufacturer |
|---|---|---|---|---|
| **CD4** | T cell | 1:500 | 30 min | Dako |
| **CD8** | T cell | 1:500 | 30min | Dako |
| **CD68** | Macrophages | 1:6000 | 30 min | Dako |
| **TIM-3** | Checkpoint | 1:500 | Over night | Abcam |
| **TIGIT** | Checkpoint | 1:200 | Over night | Abcam |
| **LAG-3/CD223** | Checkpoint | 1:300 | Over night | LSBIO |
| **ICOS/CD278** | Checkpoint | 1:200 | Over night | Abcam |
| **CTLA-4/CD152** | Checkpoint | 1:300 | Over night | Origene |
| **PD-1/CD279** | Checkpoint | 1:250 | Over night | Abcam |
| **PD-L1/CD274** | Checkpoint | 1:200 | Over night | Dako |
| **PD-L2/CD273** | Checkpoint | 1:400 | Over night | Abcam |

Supplementary table 3. Genes in Nanostring Pan Cancer Progression panel,

AAMP, BI3BP, CHE, CTG2, CVR1, CVR1C, CVRL1, DAM15, DAM17, DAM28, DAM8, DAM9, DAMTS1, DAMTS12, DAMTS8, DAP1, DD1, DM2, DRA2B, EBP1, AGGF1, AGR2, AGRN, AGT, AHNAK, AKAP12, AKAP2, AKT1, AKT2, AKT3, ALB, ALDOA, ALOX5, AMH, ANG, ANGPT1, ANGPT2, ANGPTL2, ANGPTL4, ANPEP, ANXA2P2, AP1M2, APC, APOD, APOE, APOH, AQP1, ARAP2, AREG, ARHGAP32, ARHGDIB, ASPN, ATPIF1, B3GNT3, BAD, BAG2, BAI1, BAI3, BCAS1, BGN, BICC1, BMP4, BMP5, BMP7, BMPER, BMPR1A, BMPR1B, BMPR2, BNC2, BRMS1, BTG1, C1S, C3, C3AR1, CADM1, CALCRL, CALD1, CAMK2A, CAMK2B, CAMK2D, CAMP, CASP8, CAV1, CBLC, CCBE1, CCDC80, CCL11, CCL21, CCL5, CCL7, CCL8, CCR2, CCR3, CD163, CD24, CD2AP, CD34, CD36, CD44, CD46, CD82, CDC42, CDH1, CDH11, CDH13, CDH2, CDK14, CDKN1A, CDKN2A, CDS1, CEACAM1, CEACAM5, CEACAM6, CEP170, CEP295, CFP, CGN, CHAD, CHD4, CHI3L1, CHP1, CHP2, CHRDL1, CHRNA7, CIB1, CKMT1A, CLDN1, CLDN3, CLDN4, CLDN7, CLEC2B, CLEC3B, CLIC4, CLU, CMA1, CNN1, COL18A1, COL1A1, COL1A2, COL3A1, COL4A1, COL4A2, COL4A6, COL5A1, COL5A2, COL6A1, COL6A2, COL6A3, COL7A1, COMP, CREBBP, CRIP2, CRISPLD2, CSF2RB, CSPG4, CST7, CTNNB1, CTNND1, CTSG, CTSH, CTSK, CTSL, CUL1, CX3CL1, CXADR, CXCL10, CXCL11, CXCL12, CXCL13, CXCL17, CXCL8, CXCR2, CXCR3, CXCR4, CYB561, CYBB, CYP1B1, DAG1, DCC, DCN, DDR2, DENND5A, DENR, DESI1, DICER1, DLC1, DLG1, DLL4, DPT, DPYSL3, DSC2, DST, ECM1, ECM2, ECSCR, EDN1, EGF, EGFL7, EGFR, EGLN2, EGLN3, EIF2AK3, EIF4E2, EIF4EBP1, ELF3, ELK3, EMCN, EMILIN1, EMILIN3, EMP3, ENO1, ENO2, ENO3, ENPEP, ENPP2, EP300, EPAS1, EPCAM, EPHA1, EPHA2, EPHB1, EPHB3, EPHB4, EPN3, EPS8L1, ERBB2, ERBB2IP, ERBB3, EREG, ERMP1, ESRP1, ETV4, EVI2A, EVPL, F11R, F3, FAM174B, FAP, FASLG, FBLN1, FBLN5, FBN1, FBN2, FBP1, FERMT2, FGF18, FGF2, FGF9, FGFR1, FGFR2, FGFR3, FGFR4, FGL2, FHL1, FIGF, FLI1, FLT1, FLT4, FMOD, FN1, FOXC2, FOXO4, FRAS1, FREM1, FREM2, FST, FSTL1, FUT3, FXYD6, GALNT7, GATA4, GDF15, GDF5, GDF6, GIMAP4, GIMAP6, GJA5, GLYR1, GPI, GPR124, GPR56, GPX1, GREM1, GRHL2, GSN, GTF2I, GZMK, HAPLN1, HAS1, HDAC5, HDHD3, HEG1, HGF, HIF1A, HIPK1, HIPK2, HK2, HK3, HKDC1, HLA-DPB1, HMOX1, HOXA5, HOXA7, HOXB13, HOXB3, HPSE, HRAS, HSD17B12, HSP90B1, HSPB1, HSPG2, HUNK, IBSP, ICAM1, ID1, ID2, ID4, IFNG, IGF1, IGFBP4, IGFBP7, IL10RA, IL11, IL13RA2, IL15, IL18, IL1A, IL1B, IL1RL1, IL1RN, IL6, ILK, INHBA, INHBE, IRF6, ISL1, ISLR, ITGA1, ITGA11, ITGA2, ITGA3, ITGA5, ITGA6, ITGA7, ITGA8, ITGA9, ITGAM, ITGB1, ITGB1BP1, ITGB2, ITGB3, ITGB4, ITGB6, ITGB7, ITGB8, ITM2A, JAG1, JAM2, JAM3, JUN, KCNJ8, KDM1A, KDR, KIAA1462, KISS1, KLK3, KRAS, KRIT1, KRT1, KRT14, KRT19, KRT7, LAD1, LAMA1, LAMA3, LAMA4, LAMA5, LAMB3, LAMC1, LAMC2, LDHA, LEFTY1, LGALS1, LHFP, LIFR, LLGL2, LOX, LOXL2, LRG1, LTBP4, LUM, LY96, MAF, MAP2K1, MAP2K2, MAP2K4, MAP3K7, MAPK1, MAPK3, MAPKAPK3, MCAM, MED1, MED23, MEG3, MEOX2, MET, MFAP4, MGAT5, MGP, MISP, MMP1, MMP10, MMP12, MMP13, MMP14, MMP17, MMP2, MMP24, MMP3, MMP9, MMRN2, MPDZ, MRC1, MS4A4A, MS4A6A, MT3, MTA1, MTBP, MTDH, MTOR, MUC1, MYC, MYCL, MYH11, MYLK, MYO1D, MYO5C, NAA15, NAP1L3, NCAM1, NCL, NDNF, NDP, NDRG1, NF1, NF2, NFAT5, NFATC2, NFKB1, NID2, NME1, NME4, NODAL, NOS2, NOS3, NOTCH1, NOX5, NPR1, NR3C1, NR4A1, NR4A3, NRCAM, NRP1, NRP2, NRXN1, NRXN3, NTRK1, OAS1, OCLN, OGN, OLFML2B, OVOL2, P3H1, P3H2, PCOLCE, PDCD10, PDCL3, PDGFA, PDGFC, PDGFRB, PDK1, PDPN, PEBP4, PECAM1, PFKFB1, PFKFB4, PGK1, PIK3CA, PIK3CD, PIK3CG, PIK3R1, PIK3R2, PIK3R5, PIK3R6, PITX2, PKM, PKN1, PKNOX1, PLA2G10, PLA2G2A, PLA2G2D, PLA2G3, PLAU, PLAUR, PLCG1, PLCG2, PLEKHO1, PLS1, PLXDC1, PLXNC1, PLXND1, PMP22, PNPLA6, POPDC3, POSTN, PPFIBP2, PPL, PPP1R16B, PPP2CB, PPP2R1A, PPP3R1, PRELP, PRF1, PRKCB, PRKCG, PRKCZ, PROK2, PROM1, PRR15L, PRSS22, PRSS8, PTEN, PTGDS, PTGIS, PTGS2, PTK2, PTK2B, PTK6, PTPRB, PTPRC, PTPRM, PTRF, PTTG1, PTX3, PXDN, PYCARD, QKI, RAB25, RAC1, RAC2, RAF1, RAMP1, RAMP2, RB1, RBL1, RBL2, RBM47, RBPJ, RBX1, RELN, RGCC, RHOA, RNH1, ROBO4, ROCK1, ROCK2, RORA, RORB, RPS27A, RPS6KB1, RPS6KB2, RRAS, RTN4, RUNX1, RUNX1T1, S100A14, S100A7, S1PR1, SACS, SAMSN1, SCG2, SCNN1A, SDC4, SELE, SEMA3E, SERINC5, SERPINA1, SERPINE1, SERPINF1, SERPING1, SERPINH1, SET, SETD2, SFRP1, SFRP2, SH2B3, SH2D3A, SH3YL1, SHB, SIRT1, SKP1, SLC12A6, SLC2A1, SLC35A3, SLC37A1, SLC44A4, SLIT2, SLPI, SMAD1, SMAD2, SMAD3, SMAD4, SMAD5, SMAD9, SMC3, SMOC1, SMURF1, SMURF2, SNAI1, SNAI2, SNAI3, SNRPF, SOD1, SORD, SOX17, SOX2, SOX9, SP1, SPARC, SPARCL1, SPDEF, SPHK2, SPINK5, SPINT1, SPOCK3, SPP1, SRC, SRF, SRGN, SRPK2, SRPX2, SSTR2, ST14, STAB1, STAB2, STAT1, STAT3, SULF1, SV2B, SYK, SYNE1, TACSTD2, TAL1, TBX1, TBX4, TBXA2R, TCEB1, TCEB2, TCF20, TCF3, TCF4, TDGF1, TEK, TF, TFDP1, TFPI2, TGFB1, TGFB2, TGFBI, TGFBR2, THBS1, THBS2, THBS4, THY1, TIE1, TIMP1, TIMP2, TIMP4, TJP2, TJP3, TLR4, TMC6, TMEM100, TMEM30B, TMPRSS2, TMPRSS4, TMPRSS6, TNC, TNF, TNFRSF12A, TNFRSF1A, TNFSF10, TNFSF12, TNFSF13, TNMD, TNN, TNS1, TNXB, TOM1L1, TP53, TPM2, TPSB2, TPSD1, TSHR, TSPAN1, TWIST1, TWIST2, TXNIP, TYMP, UBA52, UTS2, VAMP8, VASH1, VAV2, VAV3, VCAM1, VCAN, VEGFA, VEGFB, VEGFC, VEZF1, VHL, VIM, VIT, VPS13A, VSIG4, VWA1, VWA2, WARS, WIPF1, WNT5A, WNT5B, WWTR1, ZC3H12A, ZCCHC24, ZEB1, ZEB2, ZFPM2, ZFYVE16, ZFYVE9, AGK, AMMECR1L, CC2D1B, CNOT10, CNOT4, COG7, DDX50, DHX16, DNAJC14, EDC3, EIF2B4, ERCC3, FCF1, GPATCH3, HDAC3, MRPS5, MTMR14, NOL7, NUBP1, PRPF38A, SAP130, SF3A3, TLK2, TMUB2, TRIM39, USP39, ZC3H14, ZKSCAN5, ZNF143, ZNF346

Supplementary table 4. Patient characteristics validation cohort

| | | |
|---|---|---|
| **Included pCCA patients** | 37 | |
| **Mean age (years)** | 66.16 | |
| **Mean overall survival (months)** | 13. 47 | |
| **Gender** | | |
| %Male | 22 | 59% |
| %Female | 19 | 41% |
| **Tumor stadium (T)** | | |
| %UICC T1 | 2 | 5% |
| %UICC T2 | 23 | 62% |
| %UICC T3 | 8 | 22% |
| %UICC T4 | 4 | 11% |
| **Nodal status (N)** | | |
| %N0 | 21 | 57% |
| %N1+N2 | 16 | 43% |
| **Perineural Invasion (Pn)** | | |
| %Pn0 | 7 | 19% |
| %Pn1 | 28 | 76& |
| %not known | 2 | 5% |
| **Residual Tumor (R)** | | |
| %R0 | 32 | 86% |
| %R1 | 5 | 14% |
| **Lymphovascular invasion (L)** | | |
| %L0 | | |
| %L1 | 30 | 81% |
| %not known | 7 | 19% |

Supplementary table 5. Statistical changes in marker expression original cohort

| Marker | ROI 1 | ROI 2 | n 1 | n 2 | P-value | Sign. Level | Log2 FoldChange |
|---|---|---|---|---|---|---|---|
| CD8 | T | PT | 38 | 28 | 0.046 | * | 0.376171 |
| CD68 | T | PT | 38 | 28 | 8.75E-04 | *** | 1.107765 |
| CD4_CTLA4 | T | IM | 36 | 12 | 1.62E-05 | **** | 5.642751 |
| CD4_CTLA4 | T | PT | 36 | 25 | 3.41E-06 | **** | 3.646778 |
| CD4_ICOS | T | IM | 36 | 12 | 3.47E-05 | **** | 5.399441 |
| CD4_ICOS | T | PT | 36 | 25 | 4.23E-04 | *** | 3.007225 |
| CD4_ICOS_CTLA4 | T | IM | 36 | 12 | 2.19E-05 | **** | 6.552339 |
| CD4_ICOS_CTLA4 | T | PT | 36 | 25 | 2.62E-05 | **** | 3.391128 |
| CD4_ICOS_TIGIT_CTLA4 | T | IM | 36 | 12 | 3.29E-05 | **** | 9.617247 |
| CD4_ICOS_TIGIT_CTLA4 | T | PT | 36 | 25 | 5.99E-06 | **** | 5.346448 |
| CD4_PD1_LAG-3_TIM3 | T | PT | 39 | 27 | 0.034 | * | 4.633537 |
| CD4_PD1_TIM3 | T | PT | 39 | 27 | 0.03 | * | 4.148948 |
| CD4_TIGIT | T | IM | 36 | 12 | 1.30E-05 | **** | 6.827088 |
| CD4_TIGIT | T | PT | 36 | 25 | 8.36E-06 | **** | 3.746301 |
| CD4_TIGIT_CTLA4 | T | IM | 36 | 12 | 4.07E-05 | **** | 7.048764 |
| CD4_TIGIT_CTLA4 | T | PT | 36 | 25 | 1.53E-06 | **** | 5.227363 |
| CD4_TIM3 | T | PT | 39 | 27 | 0.04 | * | 1.396392 |
| CD68_PD1 | T | PT | 38 | 28 | 0.01 | ** | 0.681096 |
| CD68_PDL1 | T | PT | 38 | 28 | 0.009 | ** | 0.832505 |
| CD68_PDL2 | T | PT | 38 | 28 | 1.63E-04 | *** | 1.013609 |
| CD8_CTLA4 | T | IM | 36 | 12 | 0.001 | *** | 2.328888 |
| CD8_CTLA4 | T | PT | 36 | 25 | 4.95E-07 | **** | 2.550666 |
| CD8_ICOS | T | IM | 36 | 12 | 4.87E-05 | **** | 3.559524 |
| CD8_ICOS | T | PT | 36 | 25 | 4.62E-06 | **** | 2.878366 |
| CD8_ICOS_TIGIT | T | IM | 36 | 12 | 3.49E-06 | **** | 8.762447 |
| CD8_ICOS_TIGIT | T | PT | 36 | 25 | 5.95E-06 | **** | 4.183281 |
| CD8_ICOS_TIGIT_CTLA4 | T | IM | 36 | 12 | 4.62E-05 | **** | 9.464115 |
| CD8_ICOS_TIGIT_CTLA4 | T | PT | 36 | 25 | 6.50E-06 | **** | 4.25536 |
| CD8_LAG-3 | T | PT | 39 | 27 | 0.018 | * | 0.777919 |
| CD8_LAG-3_TIM3 | T | PT | 39 | 27 | 0.01 | ** | 1.911776 |
| CD8_PD1_LAG-3 | T | PT | 39 | 27 | 0.011 | * | 3.03886 |
| CD8_PD1_LAG-3 | IM | PT | 5 | 27 | 0.037 | * | 1.040324 |
| CD8_PD1_LAG-3_TIM3 | T | PT | 39 | 27 | 0.003 | ** | 4.827998 |
| CD8_PD1_TIM3 | T | PT | 39 | 27 | 8.08E-04 | *** | 4.497732 |
| CD8_TIGIT | T | IM | 36 | 12 | 4.79E-05 | **** | 5.05979 |
| CD8_TIGIT | T | PT | 36 | 25 | 7.87E-06 | **** | 2.360652 |
| CD8_TIGIT_CTLA4 | T | IM | 36 | 12 | 1.04E-04 | *** | 5.920656 |
| CD8_TIGIT_CTLA4 | T | PT | 36 | 25 | 2.69E-06 | **** | 3.509071 |
| CD8_TIM3 | T | PT | 39 | 27 | 0.002 | ** | 0.141312 |

Combination of markers with significantly increased or decreased percentage of positive cells in one of the ROIs (CT = central tumor, PT = peri-tumor, IM = Invasive margin). Shown are only comparisons with a p-value < 0.05. n1 and n2 indicate the number of patients/ slides with data for the marker in each ROI. P values were generated using Mann-Whitney U test and Fold change is shown as a log2. * p < 0.05, **p < 0.01, ***: p < 0.001, **** p <= 0.0001.

Supplementary table 6. Statistical changes in marker expression validation cohort

| Marker | ROI 1 | ROI 2 | n 1 | n 2 | P-value | Sign. Level | Log2 FoldChange |
|---|---|---|---|---|---|---|---|
| CD4_LAG-3_TIGIT | T | PT | 24 | 5 | 0.006 | ** | 5.676495 |
| CD4_LAG-3_TIGIT_CTLA44 | T | PT | 24 | 5 | 0.002 | ** | 6.410646 |
| CD4_TIGIT | T | PT | 24 | 5 | 0.006 | ** | 5.660531 |
| CD8 | T | IM | 24 | 5 | 7.07E-04 | *** | -1.59012 |
| CD8 | IM | PT | 5 | 5 | 0.032 | * | 1.993647 |
| CD8_CTLA44 | T | PT | 24 | 5 | 0.046 | * | -0.12361 |
| CD8_LAG-3 | T | IM | 24 | 5 | 2.02E-04 | *** | -1.99452 |
| CD8_LAG-3 | IM | PT | 5 | 5 | 0.016 | * | 1.949832 |
| CD8_LAG-3_CTLA44 | T | PT | 24 | 5 | 0.046 | * | -0.21226 |
| CD8_LAG-3_CTLA44_TIGIT | T | PT | 24 | 5 | 8.88E-04 | *** | 8.919423 |
| CD8_TIGIT | T | PT | 24 | 5 | 7.07E-04 | *** | 8.134363 |

, Combination of markers with significantly increased or decreased percentage of positive cells in one of the ROIs (CT = central tumor, PT = peri-tumor, IM = invasive margin). Shown are only comparisons with a p-value < 0.05. n1 and n2 indicate the number of patients/ slides with data for the marker in each ROI. P values were generated using Mann-Whitney U test and Fold change is shown as a log2. * p < 0.05, **p < 0.01, ***: p < 0.001, **** p <= 0.0001.

Supplementary table 7. Differentially expressed genes in TIGIT high patients

| **Downregulated in TIGIT high** | | | |
|---|---|---|---|
| **Gene** | **Log2 fold change** | **Linear fold change** | **P-value** |
| RHOA | -0.517 | 0.699 | 0.00248 |
| HPSE | -0.623 | 0.649 | 0.00643 |
| PLAUR | -1.07 | 0.477 | 0.00798 |
| SET | -0.522 | 0.696 | 0.0102 |
| ACHE | -1.63 | 0.324 | 0.0109 |
| LUM | -0.568 | 0.675 | 0.0115 |
| CTSK | -0.824 | 0.565 | 0.0128 |
| SLC2A1 | -1.11 | 0.462 | 0.0131 |
| GPX1 | -0.646 | 0.639 | 0.0137 |
| GALNT7 | -0.899 | 0.536 | 0.0146 |
| COMP | -1.15 | 0.45 | 0.0194 |
| TNFRSF12A | -0.831 | 0.562 | 0.0206 |
| WARS | -0.702 | 0.615 | 0.0233 |
| ANXA2P2 | -0.77 | 0.586 | 0.0242 |
| POSTN | -1.41 | 0.376 | 0.0259 |
| COL6A3 | -0.847 | 0.556 | 0.0265 |
| PTTG1 | -0.656 | 0.635 | 0.0275 |
| PKM | -0.579 | 0.669 | 0.0328 |
| TACSTD2 | -1.16 | 0.447 | 0.0339 |
| CCL11 | -0.743 | 0.598 | 0.036 |
| MMP2 | -0.703 | 0.614 | 0.0364 |
| CDKN1A | -0.549 | 0.684 | 0.0393 |
| GREM1 | -0.952 | 0.517 | 0.0399 |
| PLAU | -1.03 | 0.489 | 0.04 |
| MMP14 | -0.697 | 0.617 | 0.041 |
| TNFSF13 | -1.19 | 0.437 | 0.0438 |
| CLDN7 | -0.838 | 0.559 | 0.0454 |
| SLPI | -0.858 | 0.552 | 0.0455 |
| **Upregulated in TIGIT high** | | | |
| **Gene** | **Log2 fold change** | **Linear fold change** | **P-value** |
| PTPRB | 0.59 | 1.51 | 0.0459 |
| SEMA3E | 2.57 | 5.96 | 0.0109 |

Differential expression (DE) gene analysis was performed between the TIGIT high and TIGIT low group. Genes with Log2 Fold change > 0.5/ < - 0.5 and a p-value < 0.05 were considered DE genes. Genes that were determined to be outliers were removed using Tukey’s rule.
